# Growth of *Corynebacterium glutamicum* on 5-oxo-L-proline (pyroglutamate) as carbon and nitrogen source requires the PxpR-controlled *pxpTABC* genes

**DOI:** 10.1101/2025.10.30.685629

**Authors:** Lea Sundermeyer, Christina Mack, Michael Bott

## Abstract

5-Oxo-L-proline (5-OP) is inevitably formed in all cells by spontaneous cyclization of L-glutamate, L-glutamine, or γ-glutamyl phosphate. Its use as a substrate by bacteria has rarely been described. Here, we show that the actinobacterial species *Corynebacterium glutamicum* can grow well in minimal medium with 5-OP as sole carbon and nitrogen source. We identified the *pxpTABC* gene cluster as being essential for growth on 5-OP. The *pxpT* gene encodes a secondary transporter of the APC superfamily which most likely catalyzes 5-OP uptake into the cell. The *pxpABC* genes encode a recently identified ATP-dependent 5-oxoprolinase that converts 5-OP into L-glutamate for further metabolism. Both PxpT and PxpABC were required for growth with 5-OP. Upstream and divergent to *pxpT*, the gene *pxpR* is located, encoding a GntR-type transcriptional regulator. Deletion of *pxpR* improved growth on 5-OP suggesting that PxpR acts as repressor of the *pxpTABC* operon. This function was further supported by reporter gene studies. Purified PxpR was shown by isothermal titration calorimetry (ITC) to bind 5-OP with a K_D_ of 726 ± 23 nM. L-proline, L-glutamate, L-glutamine, and L-aspartate were not bound under the conditions tested, suggesting that PxpR is a specific 5-OP biosensor.

**IMPORTANCE:** The utilization of 5-OP as a carbon and nitrogen source for bacteria has rarely been investigated in the past, although this metabolite is ubiquitous in cells due to its non-enzymatic formation from the amino acids L-glutamate and L-glutamine. In *Corynebacterium glutamicum* cells, but also in other bacteria such as *Escherichia coli*, L-glutamate is present at concentrations in the range of 100 mM, suggesting a substantial and continuous 5-OP formation. Our results demonstrate that 5-OP can serve both as carbon and nitrogen source for *C. glutamicum* and presumably also for other bacteria.

## INTRODUCTION

5-Oxo-L-proline (5-OP), also termed pyroglutamate, is the cyclic lactam of glutamate. Its presence in all cells is inevitable due to the spontaneous cyclization of L-glutamate, L-glutamine, and γ-glutamyl phosphate (1). In the case of L-glutamine, approximately 10% is converted to 5-OP per day at 37°C with ammonia as second product (2, 3). 5-OP is also formed enzymatically in the degradation of glutathione within the γ-glutamyl cycle (4) and by cleavage of N-terminal 5-OP residues of proteins by pyroglutamyl peptidase (5, 6).

Earlier reports on 5-OP utilization and metabolism by bacteria were reviewed by van der Werf and Meister (7). A systematic study on the effects of 5-OP on different prokaryotic species revealed that growth of thermoacidophilic archaea and some bacteria was severely inhibited by the presence of 5-OP and that the extent of inhibition increased with increasing growth temperature and decreasing pH (8). For example, 15 mM 5-OP completely inhibited growth of *Sulfolobus solfataricus* at 78°C and pH 3 (9). In contrast, growth of some organisms living at neutral pH and moderate temperatures was not inhibited and sometimes even stimulated by 5-OP (8). In halophilic and alkaliphilic methanotrophs, 5-OP can serve as osmoprotectant (10). For *Salmonella enterica* serovar *typhimurium* it was reported that 5-OP cannot be used as nutrient source for growth, but has an influence on cellular aggregation (11).

5-OP formed during degradation of glutathione in eukaryotes is known to be converted to glutamate in an ATP-dependent reaction catalyzed by 5-oxoprolinase (12). An ATP-independent 5-oxoprolinase was identified in *Alcaligenes faecalis* (13). However, the equilibrium constant of the reaction, K_eq_ ([L-glutamate]/[5-oxo-L-proline]) was found to be ∼0.035, showing that the formation of 5-OP from L-glutamate is highly favored. Interestingly, sequencing of the corresponding gene revealed that the ATP-independent 5-oxoprolinase contains a Sec signal peptide and therefore is presumably active in the periplasm (14). In a recent study it was shown that of 984 analyzed bacterial and archaeal genomes only 115 contain homologs of the eukaryotic-type 5-oxoprolinase and only 10 had homologs of the enzyme of *A. faecalis* (15). The majority of the analyzed genomes was found to encode another type of ATP-dependent 5-oxoprolinase showing no sequence similarity to the eukaryotic-type enzyme. It is composed of three subunits and the corresponding genes were named *pxpA*, *pxpB* and *pxpC* for prokaryotic 5-oxoprolinase (15). This enzyme was originally identified in *Pseudomonas* (16). It is assumed that PxpB and PxpC are responsible for ATP-dependent phosphorylation of 5-OP, whereas PxpA catalyzes the decyclization to γ-glutamyl phosphate and the subsequent dephosphorylation to L-glutamate (17, 18).

Studies with *Bacillus subtilis* revealed that the *pxpABC* genes are essential for utilization of 5-OP as sole nitrogen source in minimal medium, but their absence also had a negative effect on growth with ammonia as nitrogen source (15). This suggests that the absence of 5-oxoprolinase impairs growth fitness of *B. subtilis* also in the absence of exogeneous 5-OP. It was shown that *B. subtilis* mutants lacking *pxpA*, *pxpB*, or *pxpC* accumulate 5-OP both within the cells and in the medium, whereas the wild type does not (15). Therefore, the negative effect on growth observed for these mutants might be due to 5-OP accumulation. 5-OP can be used as sole nitrogen source by *B. subtilis*, but not by mutants lacking the 5-oxoprolinase genes (15). Recently, the role of the *pxp* genes was explored in *Clostridioides difficile* (19). In contrast to *B. subtilis*, the genes showed no effect on sporulation in *C. difficile*, but were necessary for the growth-promoting effect of 5-OP observed for the wild type of this anaerobic enteric pathogen. It was suggested that the *pxpAGBC* genes facilitate the use of 5-OP as a carbon source, nitrogen source, or both a carbon and nitrogen source (19).

In this study, we analyzed the role of 5-OP in *Corynebacterium glutamicum*. This actinobacterial species is a model organism in microbial biotechnology since it is the most important industrial amino acid producer, with L-glutamate and L-lysine being produced at a scale of several million tons per year (20–23). Also L-glutamine-producing strains of *C. glutamicum* have been developed (24). Although both L-glutamate and L-glutamine spontaneously cyclize to form 5-OP, the presence and fate of this metabolite has never been studied in *C. glutamicum*. We show that *C. glutamicum* is able to grow with 5-OP as sole carbon and nitrogen source and delineate the molecular basis. 5-OP is taken up by the secondary transporter PxpT and converted to L-glutamate by the 5-oxoprolinase PxpABC. The corresponding operon *pxpTABC* is regulated by a GntR-type transcriptional regulator, which we named PxpR and which specifically binds 5-OP with high affinity.

## RESULTS

### Growth of *C. glutamicum* with 5-OP as a sole carbon and nitrogen source

5-OP is an unavoidable metabolite in all cells formed by spontaneous cyclization of L-glutamate, L-glutamine, or γ-glutamyl phosphate. Since the cytoplasmic concentration of L-glutamate in *C. glutamicum* cells under standard conditions has been reported to be in the range of 100-200 mM (25–28), it can be expected that 5-OP is constantly formed. Nevertheless, no studies were performed so far on this metabolite, which prompted us to analyze how *C. glutamicum* deals with 5-OP. Initially we tested if 5-OP can serve as carbon and nitrogen source. As shown in Fig. 1, cultivation of the wild type ATCC 13032 (WT) in modified CGXII minimal medium lacking the nitrogen sources ammonium sulfate and urea and containing 12.9 g/L (100 mM) 5-OP enabled growth, showing that 5-OP can serve as a sole carbon and nitrogen source. As a control, standard CGXII medium with 20 g/L (NH_4_)_2_SO_4_ and 5 g/L urea as nitrogen sources and 20 g/L glucose as carbon source was used. The lag phase for growth on 5-OP was long if the cells were pre-grown in glucose minimal medium, but absent if cells were pre-cultured with 5-OP, suggesting that the genes required for 5-OP utilization are inducible (Fig. 1).

**Fig. 1.**
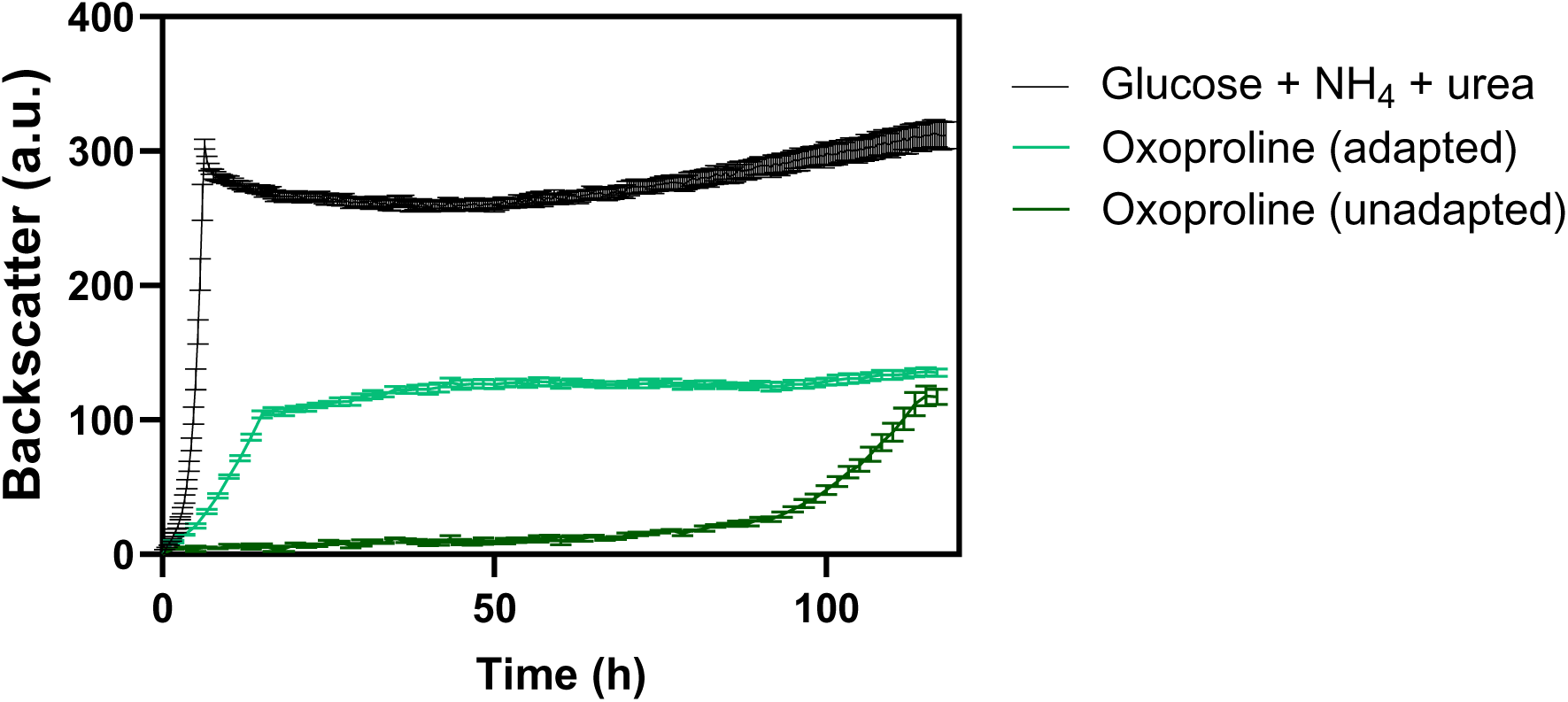
Growth of *C. glutamicum* WT on 5-OP as sole carbon and nitrogen source. Cells were cultivated either in standard CGXII medium with 20 g/L glucose as the carbon source and 20 g/L ammonium sulfate plus 5 g/L urea as nitrogen sources or in modified CGXII medium lacking glucose, ammonium sulfate, and urea and containing 12.9 g/L 5-OP as the sole carbon and nitrogen source. The pre-cultures were grown either in glucose medium (unadapted) or in 5-OP medium (adapted). Cultivation was performed in a BioLector microcultivation system at 30°C and 1200 rpm.

### Genes required for growth of *C. glutamicum* on 5-OP

Inspection of the genome of *C. glutamicum* (29, 30) for genes encoding 5-oxoprolinase revealed a cluster of three genes encoding proteins showing 31–43% amino acid sequence identity to the PxpA, PxpB, and PxpC proteins of *B. subtilis* (15). These genes were therefore named *pxpA* (cg1141)*, pxpB* (cg1140), and *pxpC* (cg1139). Upstream of *pxpA*, the gene *cg1142* is located in the same orientation (Fig. 2) that encodes a protein with 50% sequence identity to YcsG of *B. subtilis*, which is encoded downstream of *pxpA* and was shown to be required for utilization of 5-OP as a nitrogen source in *B. subtilis* (15). Based on the experiments reported below, cg1142 was named *pxpT* for 5-OP transport.

**Fig. 2.**
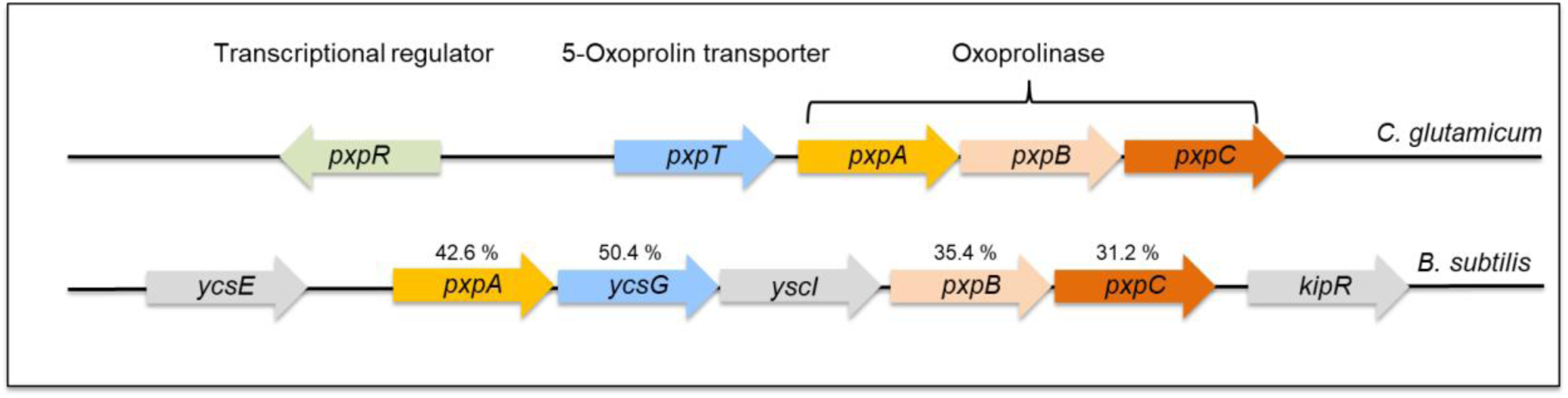
Map of the *C. glutamicum* genome region with the 5-oxoprolinase genes *pxpABC* and for comparison the corresponding region of *B. subtilis*. The %-numbers indicate the protein sequence identity to the homologous *C. glutamicum* proteins.

The relevance of the *pxpABC* genes and of *pxpT* for 5-OP utilization was tested by constructing and analyzing the *C. glutamicum* deletion mutants Δ*pxpABC*, Δ*pxpTABC*, and Δ*pxpT.* All three mutants grew like the WT in standard CGXII minimal medium with glucose (20 g/L) (Fig. 3A), but were unable to grow with 5-OP (12.9 g/L) as sole carbon and nitrogen source (Fig. 3B), suggesting that PxpABC and PxpT are essential for 5-OP utilization. For confirmation, the expression plasmids pPREx2-*pxpABC*, pPREx2-*pxpTABC*, and pPREx2-*pxpT* were constructed and transformed into the mutant strains. As shown in Fig. 3B, plasmid-based expression of *pxpABC* in the mutant strain Δ*pxpABC* not only restored growth on 5-OP, but even improved growth (reduced lag phase, increased growth rate) compared to the WT. Similarly, plasmid-based expression of either *pxpT* alone or of *pxpTABC* recovered growth of the Δ*pxpT* mutant on 5-OP, however, expression of *pxpTABC* enabled markedly better growth than *pxpT* alone (Fig. 3C). Complementation of the Δ*pxpTABC* mutant with plasmid-encoded *pxpTABC* recovered growth on 5-OP, whereas complementation with *pxpABC* alone or *pxpT* alone did not (Fig. 3D). These results confirm that both *pxpT* and *pxpABC* are required for growth of the Δ*pxpTABC* mutant on 5-OP. In some cases, the Δ*pxpTABC* mutant transformed with pPREx2-*pxpTABC* did not grow on 5-OP and plasmid resequencing showed that these strains lacked the transporter gene and part of 5-oxoprolinase-encoding sequence. The reason for this effect is unknown yet. We also tested whether pre-cultivation on 5-OP influences growth of the strain Δ*pxpT* with pPREx2-*pxpT*. As shown in Fig. 3E, precultivation on 5-OP strongly reduced the lag phase and increased the growth rate, suggesting that in this strain the *pxpABC* genes remain inducible by 5-OP.

**Fig. 3.**
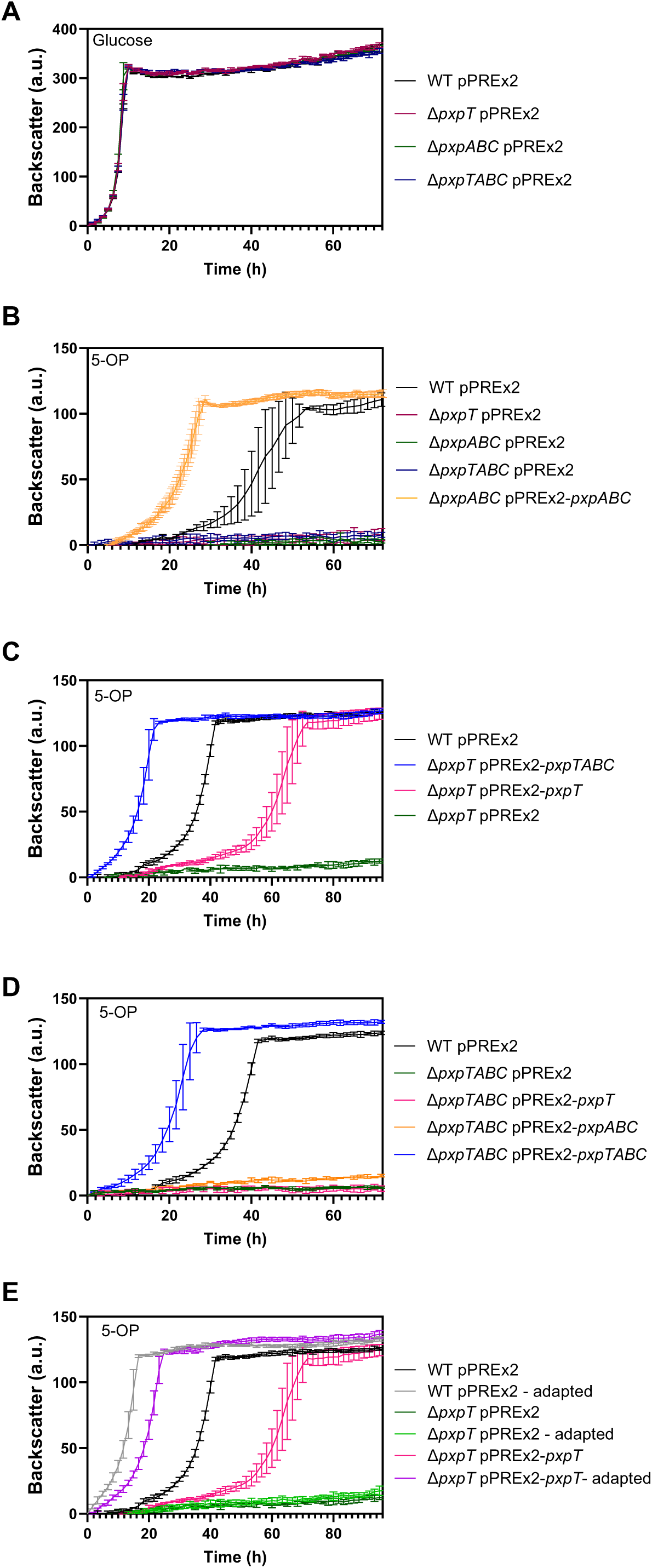
Dependency of growth with 5-OP on the *pxpTABC* genes. The indicated *C. glutamicum* strains were cultivated either in CGXII glucose medium as a control (A) or in modified CGXII medium containing 5-OP as the sole carbon and nitrogen source (B-E). All media contained 50 µg/mL kanamycin. “Adapted” indicates that the corresponding cultures were pre-grown in 5-OP medium, whereas all others were pre-grown in glucose medium. Cultivation was performed in a BioLector microcultivation system at 30°C and 1200 rpm.

### Regulation of the *pxpTABC* genes by PxpR

Upstream and divergently oriented to *pxpT*, a gene (cg1143) encoding a transcriptional regulator of the GntR family (31) is located and was designated *pxpR* (Fig. 2). PxpR consists of an N-terminal helix-turn-helix (HTH) domain (residues 11-69) and a C-terminal ligand-binding domain (residues 79-205). Due to its genomic localization, an involvement of PxpR in transcriptional regulation of the *pxpTABC* genes appeared likely and was tested by the construction and analysis of a Δ*pxpR* deletion mutant. As shown in Fig. 4, the absence of PxpR had no effect on growth on glucose, but strongly improved growth on 5-OP by shortening the lag phase and increasing the growth rate. Complementation of the Δ*pxpR* deletion mutant with plasmid pPREx2-pxpR-Strep inhibited growth on 5-OP, and this inhibition became stronger when *pxpR* expression was increased by IPTG addition. These results suggest a function of PxpR as a repressor of *pxpTABC* whose absence leads to constitutive expression of the genes.

**Fig. 4.**
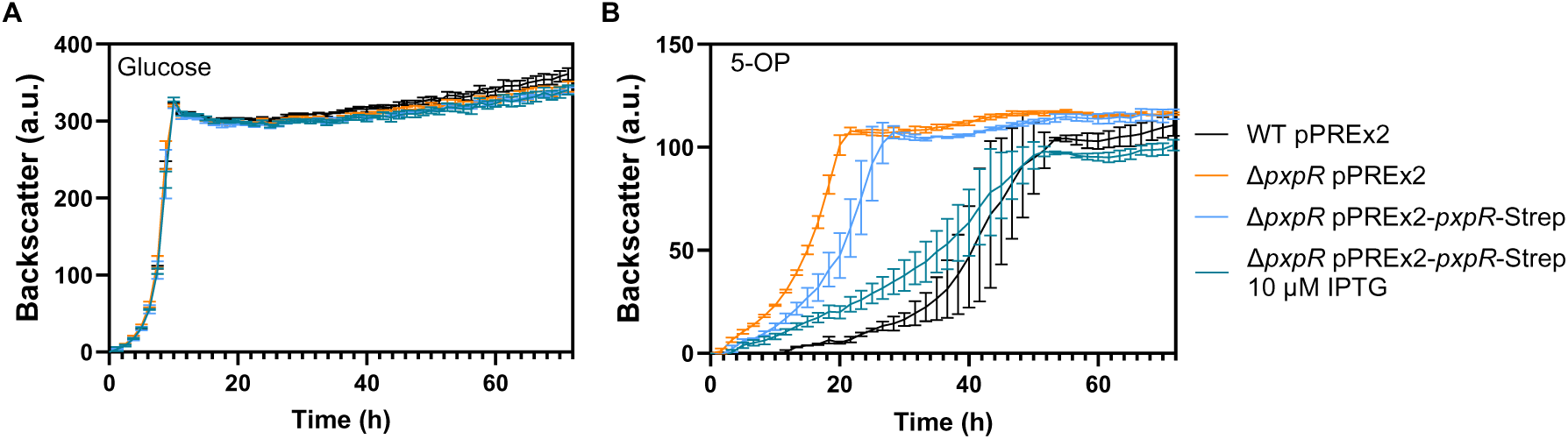
Influence of the transcriptional regulator PxpR on growth with 5-OP. The indicated *C. glutamicum* strains were cultivated either in CGXII glucose medium as control (A) or in modified CGXII medium containing 5-OP as sole carbon and nitrogen source (B), all containing 50 µg/ml kanamycin. Where indicated, the medium contained 10 µM IPTG for enhancing the expression of *pxpR*-Strep.

To confirm the repression of *pxpTABC* by PxpR, the *pxpT* promoter region (212 bp upstream of the *pxpT* start codon) was fused to the *venus* reporter gene, resulting in plasmid pJC1-P_pxpT_-Venus. The plasmid was introduced into the WT and the Δ*pxpR* mutant and the two strains were cultivated in standard CGXII glucose medium. Both strains showed the same growth behavior, but the specific Venus fluorescence (ratio of fluorescence to backscatter) was 3-fold higher for the Δ*pxpR* mutant (0.177 ± 0.005) than for the wild type (0.063 ± 0.018), supporting the role of PxpR as a repressor for the *pxpT* promoter.

### PxpR functions as a biosensor for 5-OP

Based on the results described above, it appeared likely that PxpR senses the 5-OP concentration in the cytoplasm and that binding of 5-OP to PxpR triggers a conformational change that leads to the derepression of the *pxpTABC* genes. To test for 5-OP binding to PxpR, the protein modified by a carboxyterminal Strep-tag was overproduced in *E. coli* BL21(DE3) and purified by StrepTactin affinity chromatography followed by size-exclusion chromatography (Fig. 5). The purified protein was used for binding studies using isothermal titration calorimetry, which measures binding interactions by detecting the heat absorbed or released during a binding event (32, 33). Deconvolution of the binding isotherm gives the number of binding sites (n), the enthalpy of binding (ΔH) and the equilibrium dissociation constant (K_D_). From these parameters, the Gibbs free energy (ΔG) and the entropy of binding (ΔS) can be derived. The ITC experiments (Fig. 5 and Table S1) showed that PxpR binds 5-OP with a K_D_ of 726 ± 23 nM (mean value of five determinations). Binding of 5-OP is an exothermic event with a ΔH of −98.7 ± 1.0 kJ mol^-1^ and a loss of entropy (-TΔS = 63.6 ± 1.1 kJ mol^-1^). The binding stoichiometry was 0.324 ± 0.004 if each PxpR monomer binds one molecule of 5-OP, suggesting that a fraction of the protein was in a binding-incompetent state. Importantly, L-glutamate, L-aspartate, L-glutamine, and L-proline did not bind to PxpR under the same experimental conditions, suggesting that PxpR serves as a specific 5-OP biosensor.

**Fig. 5.**
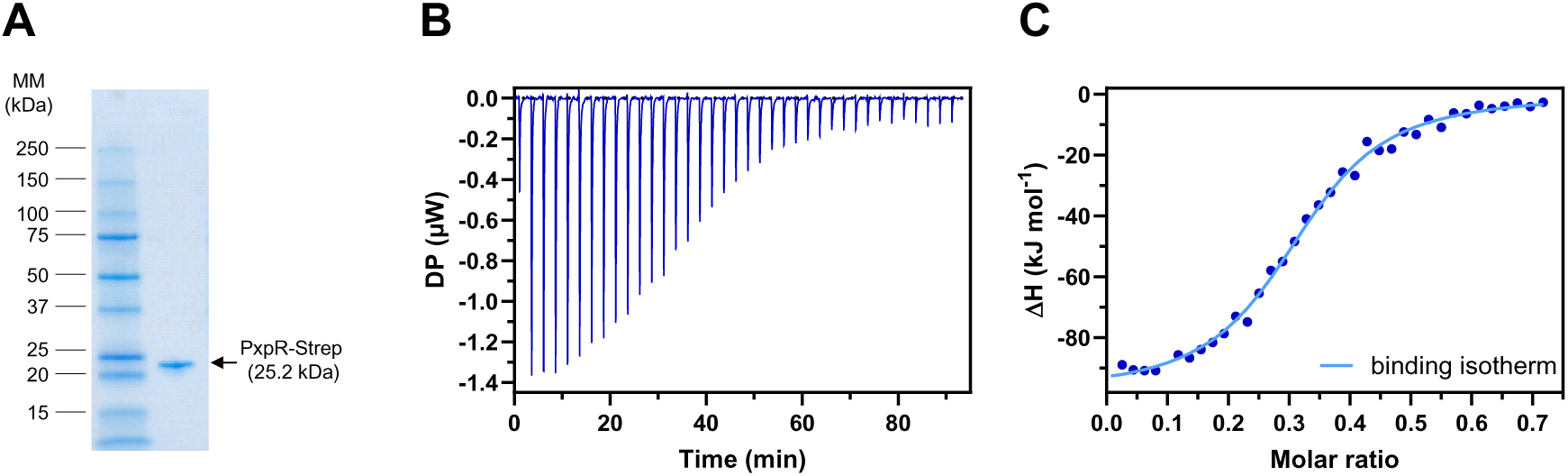
Purification of PxpR and ligand binding studies by isothermal titration calorimetry. (A) Size exclusion chromatography of PxpR-Strep purified by. (B) SDS-PAGE of PxpR-Strep purified by Strep-tag HP affinity and size exclusion chromatography. (B) Raw data for a representative ITC experiment with 40 µM PxpR-Strep and titration with 150 µM 5-OP. (C) Binding isotherm calculated from the raw data shown in Fig. 5C. DP, differential power.

## DISCUSSION

In this study we analyzed the utilization of 5-OP by *C. glutamicum*. To the best of our knowledge, no studies have been performed to date on the fate of this metabolite in *C. glutamicum*, although it is well known that 5-OP is formed spontaneously from L-glutamate, L-glutamine, and γ-glutamyl phosphate. The presence of 5-OP in *C. glutamicum* cells was previously reported in a metabolomics study by GC-MS (34). We showed that *C. glutamicum* can grow with 5-OP as the sole carbon and nitrogen source and identified the genes required for this metabolic pathway and their transcriptional regulation. The assumed pathways for 5-OP formation and utilization are summarized in Fig. 6. Due to the widespread occurrence of the *pxpABC* genes (15), it can be assumed that the capability to utilize 5-OP as a carbon and nitrogen source is widespread in bacteria.

**Fig. 6.**
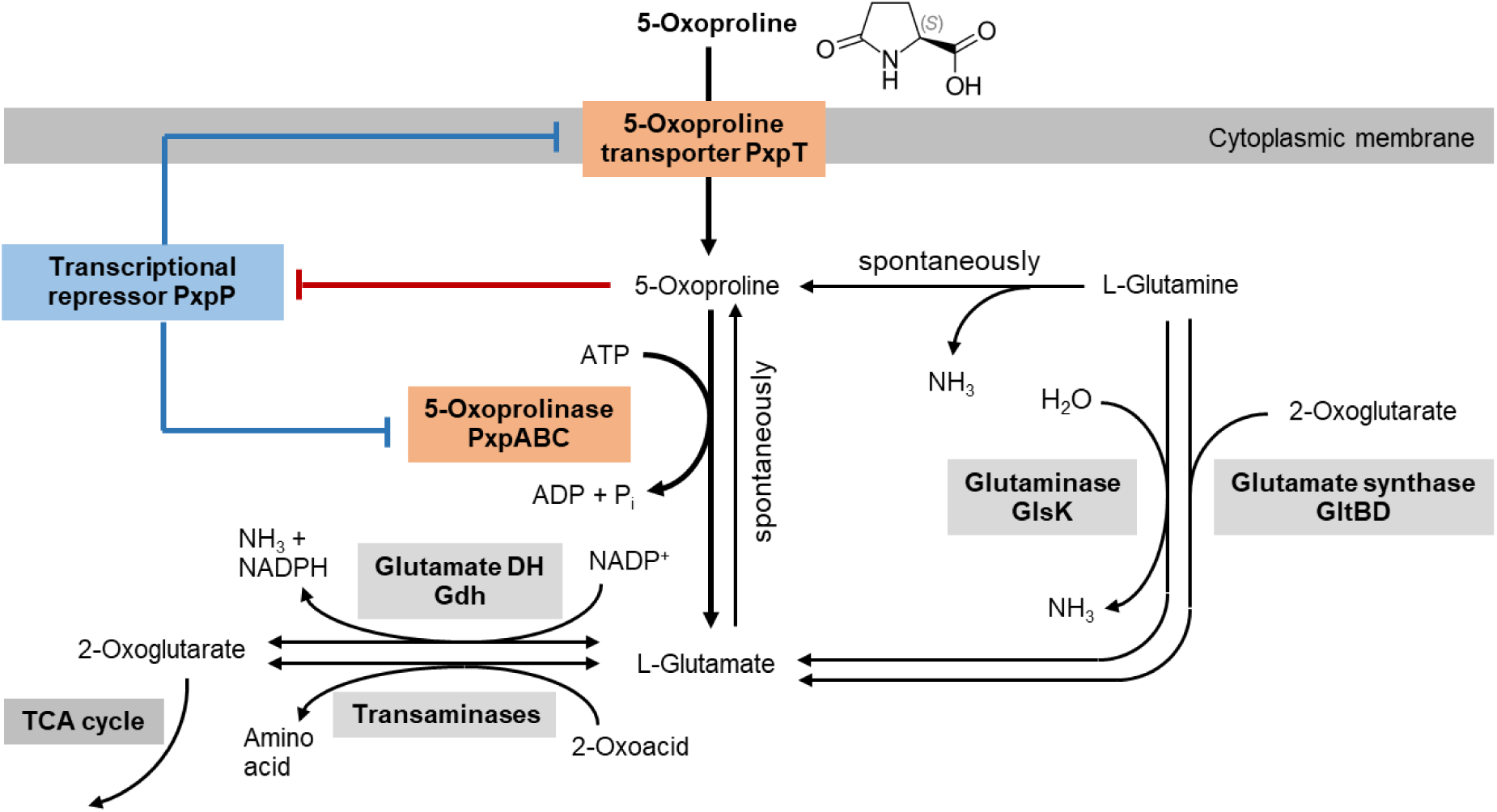
Overview on the formation and degradation of 5-OP in *C. glutamicum* and transcriptional regulation by PxpR.

The *pxpT* gene encodes a secondary transporter of 436 amino acid residues with 11 predicted transmembrane helices. BLAST searches in the Transporter Classification Database (35) showed that the proteins with the highest sequence identity, such as YcsG of *B. subtilis*, belong to the NRAMP family of divalent metal ions transporters (TCDB 2.A.55). This family is part of the large amino acid-polyamine-organoCation (APC) superfamily of secondary transporters, which also includes several families involved in amino acid transport (36). Although we cannot exclude a function of PxpT in the transport of divalent metal ions, our data suggest that PxpT is required for uptake of 5-OP by *C. glutamicum*. Deletion of *pxpT* prevented growth on 5-OP, which could be reversed by transformation of the Δ*pxpT* mutant with plasmid-encoded *pxpT*. The homologous YcsG protein of *B. subtilis* is encoded downstream of the *pxpA* gene and deletion of *ycsG* prevented the use of 5-OP as nitrogen source (15), supporting a function of YcsG in 5-OP uptake. A homolog of PxpT is also encoded in the recently analyzed *pxpAGBC* operon of *C. difficile* (PxpG, 46% amino acid sequence identity to PxpT), but its function has not been studied yet (19). The observation that lack of PxpT prevents 5-OP utilization suggests that the transporters involved in L-proline uptake in *C. glutamicum*, PutP (37), ProP, and EctP (38) cannot compensate for the absence of PxpT under the conditions used in our experiments. A different type of 5-OP uptake system has been described for *Bordetella pertussis*, which belongs to the tripartite ATP-independent periplasmic (TRAP) transport systems (39). The crystal structures of two extracytoplasmic solute receptors DctP6 and DctP7 were solved in complex with 5-OP. The K_D_ of DctP7 to 5-OP was found to be 0.3 µM. Although 5-OP utilization could not be demonstrated, the fact that the genes BP1887 and BP1891 for DctP6 and DcdP7, respectively, are located in the vicinity of a gene (locus tag BP1985) encoding a eukaryotic-type 5-oxoprolinase supports the existence of TRAP-type 5-OP uptake systems in some bacteria.

Similar to the *pxpT* gene, the *pxpABC* genes, encoding 5-oxoprolinase, were essential for growth of *C. glutamicum* on 5-OP as sole carbon and nitrogen source. Interestingly, growth of mutants lacking *pxpABC* or *pxpTABC* on glucose was not impaired (Fig. 3A), although 5-OP can presumably no longer be converted to glutamate by 5-oxoprolinase and therefore might accumulate within the cells. As L-proline is known to serve as a compatible solute in *C. glutamicum* that can accumulate to very high concentrations (40), also 5-OP may be tolerated at high concentrations. As mentioned before, 5-OP serves as a compatible solute in halophilic and alkaliphilic methanotrophic bacteria, which contain high concentrations under hyperosmotic stress (10). In *Methylomicrobium alcaliphilum* cells grown at a NaCl concentration of 1 M the concentration of 5-OP reached 0.4 M (41). Therefore, it seems possible that also other bacteria like *C. glutamicum* can tolerate high 5-OP concentrations.

To our knowledge, regulation of 5-OP metabolism in bacteria has not been studied before. We demonstrated that the *pxpTABC* genes of *C. glutamicum* are repressed by the GntR-type transcriptional regulator PxpR. Deletion of *pxpR* had no influence on growth with glucose, but strongly improved growth on 5-OP by reducing the lag phase and increasing the growth rate. This phenotype of the Δ*pxpR* mutant could be reversed by plasmid-encoded expression of *pxpR* whereby stronger expression led to stronger growth inhibition (Fig. 4). The repression of the *pxpTABC* genes by PxpR was supported by a plasmid-encoded reporter gene fusion of the *pxpT* promoter with *venus*, which revealed a threefold higher expression level in the Δ*pxpR* mutant than in the WT after growth on glucose.

Regarding the DNA binding site of PxpR, the RegPrecise database (42), which predicts transcriptional regulons based on comparative genomics (42), anticipates a DNA binding site with a 17 bp inverted repeat sequence motif shown in Fig. S1 for PxpR (Cg1143). This motif is centered 65 bp upstream of the start codon of *pxpT* (Fig. S2). A putative −10 region (TAACCT) for *pxpT* was identified 95.5 bp upstream of the *pxpT* start codon, supporting a a repressor function of PxpR. The transcriptional start site of *pxpR* was identified in a genome-wide study at the ATG start codon of pxpR (43), indicating that *pxpR* is a leaderless gene (44). Due to the 143-bp distance of the predicted PxpR binding site to the transcriptional start site of *pxpR*, autoregulation of *pxpR* expression appears unlikely. Although 5-OP can serve as sole nitrogen source, the *pxpTABC* genes were not found to be part of the nitrogen starvation stimulon, which includes the regulon of the master regulator of the nitrogen starvation response, AmtR (45–48). However, a binding site (TGTGCTATAGGACACA) for the cAMP-responsive global regulator GlxR was identified in the intergenic region between *pxpR* and *pxpT* close to the assumed −10 region of *pxpT* (49, 50), suggesting a negative influence on *pxpTABC* expression.

The function as a repressor of *pxpTABC* suggested that PxpR senses the cytoplasmic concentration of 5-OP and relieves repression of *pxpTABC* once the 5-OP level reaches a certain level. In agreement with this suggestion we could show by ITC that purified PxpR binds 5-OP with a K_D_ of 726 nM. This suggests that 5-OP concentrations in the µM range are sufficient for derepression of the *pxpTABC* genes. Since we could not detect binding of L-glutamate, L-aspartate, L-glutamine, and L-proline under the tested conditions, PxpR is presumably a specific 5-OP sensor. This offers the opportunity to develop various types of 5-OP biosensors enabling cytoplasmic 5-OP detection, e.g. based on expression of a reporter gene for a fluorescent protein under the control of the *pxpT* promoter (51) or FRET-based biosensors (52).

## MATERIALS AND METHODS

### Bacterial strains, media, and culture conditions

All bacterial strains and plasmids used in this work are listed in Table 2. *Escherichia coli* cells were cultivated at 37 °C in lysogeny broth (LB) (53) or terrific broth (TB) (12 g L^−1^ tryptone, 24 g L^−1^ yeast extract, 4 mL glycerol, 12.54 g L^−1^ K_2_HPO_4_, 2.31 g L^−1^ KH_2_PO_4_; pH 7.0) or on LB agar plates (Carl Roth, Karlsruhe, Germany). *C. glutamicum* strains were cultivated at 30°C in brain heart infusion medium (BHI; Difco Laboratories, Detroit, USA) or in CGXII medium with 20 g L^−1^ glucose (54) containing 30 mg L^-1^ 3,4-dihydroxybenzoate as an iron chelator or in modified CGXII medium lacking glucose, ammonium sulfate and urea and containing 12.9 g L^−1^ 5-oxo-L-proline as the nitrogen and carbon source. Solid media were prepared by adding 15 g L^−1^ agar to these media. To maintain plasmid stability, kanamycin was added at concentrations of 25 mg L^−1^ (*C. glutamicum*) or 50 mg L^−1^ (*E. coli*).

**Table 1.** Bacterial strains and plasmids used in this study.

| Strain or plasmid | Description | Reference or source |
| --- | --- | --- |
| ***C. glutamicum* strains** |  |  |
| ATCC 13032 | Biotin-auxotrophic wild-type strain | DSMZ |
| $\Delta pxpABC$ | Wild type derivative with in-frame deletion of *pxpABC* (cg1141-cg1140-cg1139) | This work |
| $\Delta pxpTABC$ | Wild type derivative with in-frame deletion of *pxpTABC* (cg1142-cg1141-cg1140-cg1139) | This work |
| $\Delta pxpT$ | Wild type derivative with in-frame deletion of *pxpT* (cg1142) | This work |
| $\Delta pxpR$ | Wild type derivative with in-frame deletion of *pxpR* (cg1143) | This work |
| ***E. coli* strains** |  |  |
| DH5α | F- *supE44* Δ*lacU169* (Φ80*lacZ*Δ*M15*) *hsdR17 recA1 endA1 gyrA96 thi-1 relA1* | (58) |
| BL21(DE3) | F- *ompT hsdSB(rB-mB-) gal dcm* (λclts857 *ind1 Sam7 nin5 lacUV5-T7 Gen 1*) | (61) |
| **Plasmids** |  |  |
| pK19mobsacB | Kan<sup>R</sup>; suicide vector for allelic exchange in *C. glutamicum*; *oriV<sub>E. coli</sub> oriT sacB* | (62) |
| pK19mobsacB-$\Delta pxpABC$ | Kan<sup>R</sup>; pK19mobsacB derivative containing a PCR product covering the upstream region of *pxpA* and the downstream region of *pxpC* | This work |
| pK19mobsacB-$\Delta pxpTABC$ | Kan<sup>R</sup>; pK19mobsacB derivative containing a PCR product covering the upstream region of *pxpT* and the downstream region of *pxpC* | This work |
| pK19mobsacB-$\Delta pxpT$ | Kan<sup>R</sup>; pK19mobsacB derivative containing a PCR product covering the upstream and downstream regions of *pxpT* | This work |
| pK19mobsacB-$\Delta pxpR$ | Kan<sup>R</sup>; pK19mobsacB derivative containing a PCR product covering the upstream and downstream region of *pxpR* | This work |
| pPREx2 | Kan<sup>R</sup>; pPBEx2 derivative (*P<sub>tacl</sub>*, *lacI<sub>q</sub>*, *oriC.g* from pBL1.; *oriE.c.* ColE1 from pUC18), with a consensus RBS (AAGGAG) for *C. glutamicum* | (63) |
| pPREx2-*pxpABC* | Kan<sup>R</sup>; pPREx2 derivative carrying the *pxpABC* genes | This work |
| pPREx2-*pxpTABC* | Kan<sup>R</sup>; pPREx2 derivative carrying the *pxpTABC* genes | This work |
| pPREx2-*pxpT* | Kan<sup>R</sup>; pPREx2 derivative carrying the *pxpT* gene | This work |
| pPREx2-*pxpR*-Strep | Kan<sup>R</sup>; pPREx2 derivative carrying the *pxpR* gene fused to a Strep tag-II encoding sequence at the 3'-end | This work |
| pPREx6 | Kan<sup>R</sup>; pPREx2 derivative (P<sub>T7</sub>, *lacI*<sup>q</sup>, *oriC.g* from pBL1.; *oriE.c.* ColE1 from pUC18), with a consensus RBS (AAGGAG) for *C. glutamicum* | (64) |
| pPREx6-*pxpR*-Strep | pPREx6 derivative carrying the *pxpR* gene fused to a Strep -tag-II encoding sequence at the 3'-end under the control of the T7 promoter | This work |
| pJC1 | Kan<sup>R</sup>; *E. coli*-*C. glutamicum* shuttle vector | (65) |
| pJC1-P<sub>*pxpT*</sub>-Venus | Kan<sup>R</sup>; pJC1 derivative containing the Venus reporter gene under the control of the *pxpT* promoter (transcriptional fusion) | This work |

**Table 2.**
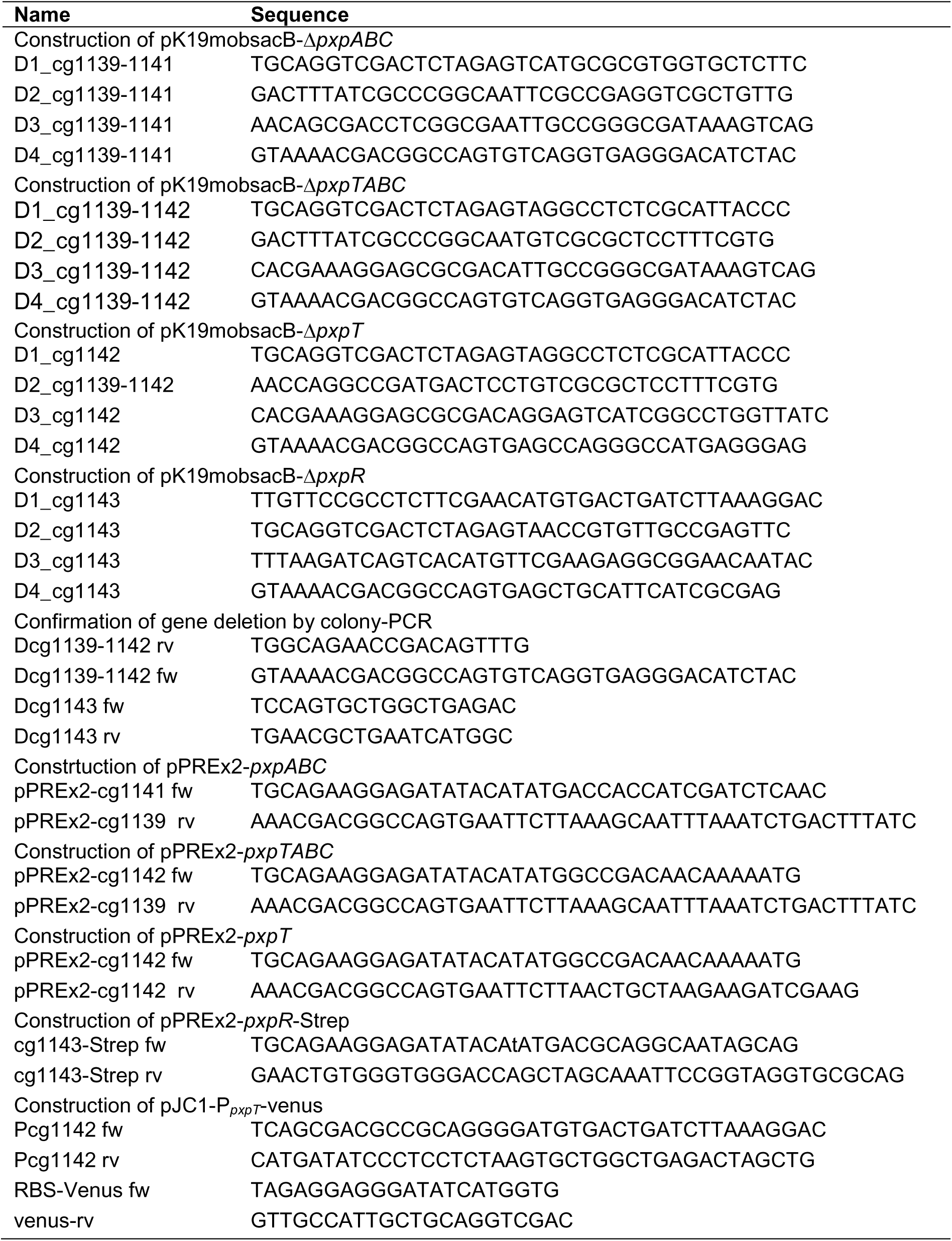
Oligonucleotides used in this study.

### Growth experiments in BioLector microcultivation systems

Growth experiments with *C. glutamicum* strains were routinely performed as microscale cultivations with BioLector I or II instruments (Beckman Coulter, Brea, USA). Growth in this system was measured online as backscattered light at 620 nm (55). Cultivations were performed in a 48-well FlowerPlate (Beckman Coulter, Brea, USA) by using the defined CGXII media described above. When strains carried expression plasmids, and 25 µg mL^−1^ kanamycin were added to the medium. Inoculations were individually prepared to reach an initial OD_600_ of 1.0. BioLector cultivations were performed at 30°C, 1200 rpm and 85% humidity. The fluorescence of the reporter protein Venus was measured with an eYFP filter (excitation 508 nm, emission 532 nm).

### Standard recombinant DNA methods and construction of deletion mutants

Standard methods such as PCR and plasmid restriction were carried out according to established protocols (56). All oligonucleotides used are listed in Table 2. Plasmids were constructed by ligating DNA fragments obtained by restriction digestion or PCR or by Gibson assembly (57). Oligonucleotide synthesis and DNA sequencing were performed by Eurofins Genomics (Ebersberg, Germany). Transformation of *E. coli* was performed using a standard protocol (58) and *C. glutamicum* transformation was performed by electroporation (59). *C. glutamicum* deletion mutants were constructed by double homologous recombination using pK19mobsacB-based plasmids as described previously (60). Oligonucleotides annealing upstream and downstream of the deleted genes were used to confirm genomic deletions by colony-PCR.

### PxpR overproduction and purification

For testing the binding of 5-OP and other metabolites to PxpR, the protein was overproduced in *E. coli* BL21(DE3) using the expression plasmid pPREx6-*pxpR*-Strep encoding a PxpR protein with a C-terminal Strep-tag-II. The strain was cultivated in TB medium with 50 µg mL^−1^ l kanamycin at 37°C. At an OD_600_ of about 0.6, *pxpR* expression was induced by addition of 0.5 mM IPTG and then the culture was further incubated overnight at 18°C. Afterwards cells were harvested and stored at −80°C. For purification of PxpR, cells were resuspended (4 g cell wet weight mL^-1^) in Strep binding buffer (100 mM Tris-HCl, 150 mM NaCl, 1 mM EDTA, pH 8) supplemented with cOMPLETE EDTA-free protease inhibitor cocktail (Roche, Basel, Switzerland) and disrupted by one passage through a Multishot Cell Disruptor (Constant System, Georgia, USA) at 20,000 psi. The cell extract was first subjected to a low-speed centrifugation (5,000 *g*, 4°C, 20 min) and subsequent ultracentrifugation of the supernatant (100,000 *g*, 4°C, 1 h). Supernatants of the ultracentrifugation were loaded onto a StrepTrap HP column (GE Healthcare, Chicago, IL, USA) and, after washing, the Strep-tagged protein was eluted using binding buffer containing 2.5 mM desthiobiotin. The protein was further purified by size exclusion chromatography on a Superdex 200 10/300 GL column (GE Healthcare, Chicago, IL, USA) equilibrated in HEPES buffer (40 mM HEPES-NaOH, 100 mM NaCl, pH 7.4). Protein concentrations were determined using a Colibri microvolume spectrometer (Berthold Detection Systems GmbH, Pforzheim, Germany) and the molar extinction coefficient at 280 nm of 22920 M^-1^ cm^-1^ predicted by the ProtParam tool (http://web.expasy.org/protparam/).

### Isothermal titration calorimetry

Purified PxpR-Strep was dialyzed overnight in HEPES buffer (40 mM HEPES–NaOH, pH 7.4, 100 mM NaCl). 20 mM stock solutions of 5-OP, L-glutamic acid, L-glutamine, L-proline, and 2-oxoglutaric acid were prepared in dialysis buffer, and the pH was adjusted to pH 7.4 using NaOH. ITC measurements were performed with a MicroCal PEAQ-ITC instrument operated at 25°C. The protein concentration was 40 μM and the ligand concentration was 200 μM. Prior to filling the measuring cell with 300 μL protein solution, the cell was rinsed with dialysis buffer, and the syringe was filled with 75 μL ligand solution. An ITC run was started with an initial injection of 0.4 μL followed by 36 injections of 1 μL each. In addition, control experiments with ligand solution titrated into the dialysis buffer were performed. The data were analyzed using the MicroCal ITC analysis software (Malvern Panalytical, Malvern, United Kingdom).

## Acknowledgements

This work was financially supported by the DFG-ANR project MetActino (DFG BO 903/4-1, ANR-18-CE92-0003). The authors thank Tobias Georgi for initial experiments and Elena von Helden for support in the preparation of the figures.

## Supplementary information to Sundermeyer et al. (2005)

**Table S1.** Thermodynamic parameters of 5-OP binding to PxpR-Strep at 25°C in 40 mM HEPES-NaOH buffer pH 7.4 containing 100 mM NaCl.

| Experiment number | PxpR (μM) | 5-OP (μM) | Binding stoichiometry | K<sub>D</sub> (nM) | ΔH (kJ mol<sup>-1</sup>) | ΔG (kJ mol<sup>-1</sup>) | -TΔS (kJ mol<sup>-1</sup>) |
| --- | --- | --- | --- | --- | --- | --- | --- |
| 1 | 40 | 150 | 0.325 | 723 | -97.7 | -35.1 | 62.6 |
| 2 | 40 | 150 | 0.322 | 746 | -98.0 | -35.0 | 63.0 |
| 3 | 40 | 150 | 0.318 | 688 | -98.1 | -35.2 | 62.8 |
| 4 | 40 | 150 | 0.330 | 742 | -99.8 | -35.0 | 64.8 |
| 5 | 40 | 150 | 0.325 | 729 | -99.7 | -35.1 | 64.6 |
| Average |  |  | 0.324 ± 0.004 | 726 ± 23 | -98.7 ± 1.0 | -35.1 ± 0.1 | 63.6 ± 1.1 |

**Fig. S1.**
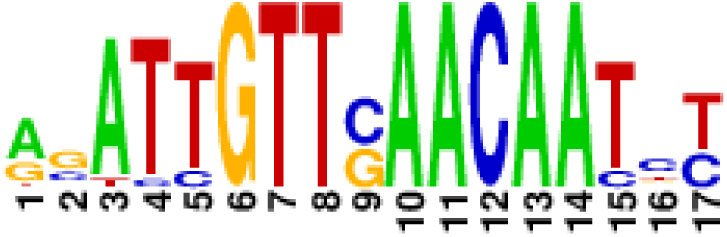
DNA-binding motif of PxpR proposed by RegPrecise (42)

**Fig. S2.**
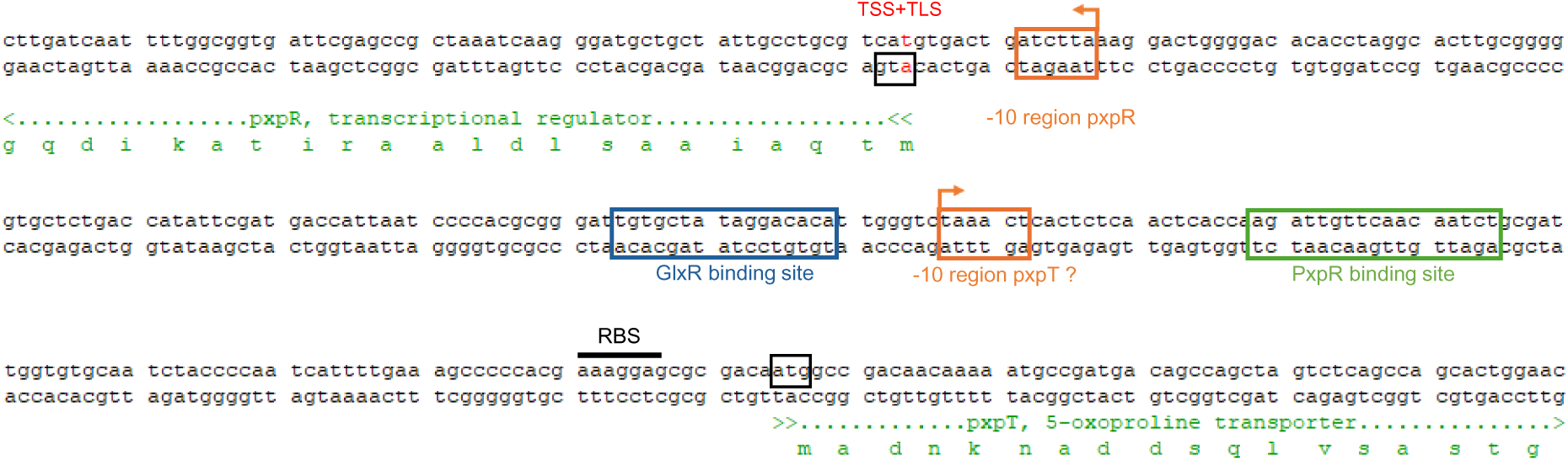
DNA sequence showing the *pxpR-pxpT* intergenic region including the initial coding sequences. The location of the putative PxpR binding site, the GlxR-binding site, and of promoter elements is indicated. The −10 region of pxpT is predicted. The *pxpR* gene is leaderless and the transcriptional start site (TSS) corresponds to the “a” nucleotide of the start codon (translational start site, TLS).

